# The Detection of *Bergeyella zoohelcum* in Young Children after Cat Bite

**DOI:** 10.1101/2021.03.26.437298

**Authors:** Min Hu, Hao Li, Tao Hu, Shuang-Yan Zhou, Ze-Hua Yang, Cai-Lan Fan, Rong Bao, Jian-Guo Shi, Ke-Bin Zhao

## Abstract

*Bergeyella zoohelcum* is an uncommon zoonotic pathogen typically associated with cat or dog bites. Previously, only 14 cases of *B. zoohelcum* infection have been reported. We isolated the bacteria from the face of a 2-year-old girl who was bitten by a cat. The organism was identified by matrix-assisted laser desorption ionization-time of flight mass spectrometry (MALDI-TOF MS) and 16S rRNA gene sequence, and phylogenetic analysise supported that this isolate was belong to *B. zoohelcum*. Due to the contradicts of culture characteristics of *B. zoohelcum* which described in reported literatures, we used different media to culture the bacteria. After 24 h incubation, Colombia blood agar (CBA), Mueller-Hinton agar plate with 5% sheep blood and chocolate agar (CA) grew well, but blood agar (BA) grew well until 72 h. The strain did not grow on McConkey agar (MAC), Mueller-Hinton agar (MHA) and chocolate agar (containing vancomycin). The low detection rate of this strain was related to its harsh growth conditions and the limitations of traditional identification techniques. With the popularization of MALDI-TOF MS and 16S rRNA gene sequence and our further understanding of the fastidious bacteria, we will quickly and accurately identify the *B. zoohelcum*, meanwhile the detection rate of this bacterium will also be significantly improved. So that clinicians can achieve precise anti-infective treatment according to antimicrobial susceptibility testing.

---

Visits to the emergency room (ER) on account of animal bites are not unusual. In fact, approximately 1% of all visits to the ER are related to bites by animals. Roughly 5 to 15% of the animal bites reported are feline inflicted (1). Cat bites are more likely to penetrate deeply, leaving only a small and deep punctures skin opening that provide access to the ideal subcutaneous breeding ground for any anaerobic opportunistic pathogen transferred from the cat’s mouth, and tend to carry a higher risk of infection and soft-tissue abscess (2). In general, the smaller the victim, the more likely a facial or scalp injury will occur; the frequency of 63% for victims less than 4 years of age are at a greatly increased risk of being bitten on the head, face, or neck (3).

Most infections caused by animal bites are polymicrobial, with mixed aerobic and anaerobic species (1). Common pathogens associated with animal bites include *Staphylococcus, Streptococcus, Pasteurella, Capnocytophaga, Moraxella, Corynebacterium, Neisseria*, and anaerobic bacteria. *Pasteurella multocida* subspecies *multocida* and *septica* were the most common isolates of cat bites (4). Up to now, a great deal of literature has commented on the pathogenic role of *Pasteurella multocida* in cat bite injuries. However, there have been few reports of *B. zoohelcum*. In our laboratory, both *Pasteurella multocida* and *B. zoohelcum* were simultaneously isolated from the secretion wound by a cat bite of a 2-year-old girl. In the literature review, only 14 cases of *B. zoohelcum* were reported, therefore, we further discussed the reasons for the low detection rate of *B. zoohelcum*.

## Case history

A healthy 2-year-old girl was bitten by a cat. Examination of the head and face showed multiple wounds, which damaged the muscle layer and was accompanied by obvious active bleeding. One hour after the girl was bitten, she was injected with rabies vaccine and tetanus immunoglobulin. In order to seek further treatment, the patient was admitted to the emergency department of hospital (The First Hospital of Shanxi Medical University, Taiyuan, shanxi, China), then, transferred to the department of plastic surgery. Laboratory evaluation revealed leukocyte count of 17,200 per cubic millimeter, with 81.5% neutrophils, 14.4% lymphocytes, and 3.7% monocytes. The girl was operated on within 2 h of admission. Basic medical management of bite wounds includes thorough cleansing and debridement. The wound was copiously irrigated with iodopor, hydrogen peroxide and normal saline. Debridement and suturing under general anesthesia was performed to remove wound edges and the tissue with poor activity. Wound specimens were taken for culture. Empirical antibiotic treatment with cefuroxime sodium 375 mg/8 h intravenously guttae was initiated. On the second day of hospitalization, all wounds healed well without further infection and swelling of her face had decreased substantially, therapy was swithed to apply mupiroxine ointment on the wounds. The patient was discharged from the hospital 8 days after admission.

## MATERIALS AND METHODS

### Bacterial strains

The strain was isolated from a 2-year-old girl after a cat bite wound.

### Culture conditions

Growth was tested on the following media: BA (Autobio, Henan, China), CBA (Autobio, Henan, China), MH agar plate with 5% sheep blood (Autobio, Henan, China), CA (Autobio, Henan, China), CA containing vancomycin (Autobio, Henan, China), MAC (Autobio, Henan, China), and MHA (Autobio, Henan, China). The plate contents were incubated at 35°C with 5% CO_2_. Growth was evaluated for the following cultured time range: 24 h, 48 h, 72 h.

### Species identification

The isolated strain was identified by Vitek 2 system (bioMérieux SA, Marcy, France) and MALDI-TOF MS (bioMérieux SA, Marcy, France).

### 16S rRNA sequence and phylogenetic analysis

The isolate was characterized by amplifying, sequencing, and analyzing the 16S rRNA gene. A fragment of approximately 1400 bp was amplified by PCR from the extracted DNA using bacterial universal primer sets 27F (5′ AGA GTT TGA TCC TGG CTC AG 3′) and 1492R (5′ TAC GGC TAC CTT GTT ACG ACT T 3′). Sequences were determined using an automatic DNA sequencer (Applied Biosystems 3730XL DNA Analyzer, Foster, USA). The 16S rRNA gene sequences were compared with the records of the GenBank database (http://www.ncbi.nlm.nih.gov/blast) using BLAST searches. A subset of the DNAMAN alignment, including the isolate 16S-D3.13302251 sequence, related sequences (representatives of the *Chryseobacterium, Cloacibacterium* and *Elizabethkingia*), and the sequence of *Weeksella virosa* ATCC 43766 (used as an outgroup), was selected for more detailed phylogenetic analysis. Sequence alignments were carried out using Clustal X. Aligned sequences were analyzed using MEGA software (MEGA version 7.0). The phylogenetic distances were calculated by the maximum-likelihood methods and a phylogenetic tree was constructed using an neighbor-joining plot program. The robustness of this tree was assessed by bootstrap method (1000 replicates with resampling of all positions).

### Antibiotic susceptibility testing (AST)

The antimicrobial susceptibility of the *B. zoohelcum* strain was determined by the Kirby-Bauer disk diffusion method on MHA plate with 5% sheep blood with use of Oxoid disk.

### Nucleotide sequence accession numbers

The GenBank accession numbers for the 16S rRNA sequence of the *B. zoohelcum* 16S-D3.13302251 strain is as follow:

MW534713.

## RESULTS

### Biochemical characteristics

The isolate grew well on the CBA, MHA plate with 5% sheep blood and CA after 24 h. While the strain did not grow on the blood agar plates after 24 h. Light growth of the strain was observed on the BA until cutrured for 48 h. The organism did not grow on CA (containing vancomycin), MAC and MHA after 24 h, 48 h and 72 h (Fig. 1). Colonies were circular, gray, wet, translucent, shiny, smooth, with entire edge and very sticky, making them difficult to remove from solid media. No hemolysis was seen on Colombia blood agar plates. The organism test positive for oxidase, catalase and indole. Urease is strongly positive immediately.

**FIG 1.**
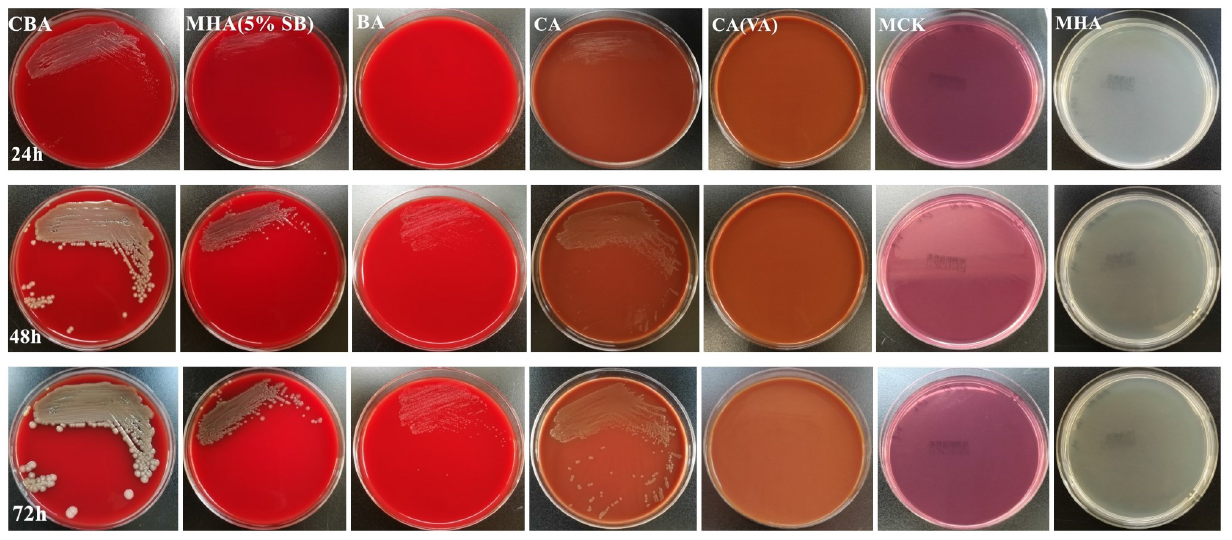
The growth of *B. zoohelcum* cultured on different media for different time

### Determination of Vitek 2 system, MALDI-TOF MS and 16S rRNA gene sequence

A Vitek 2 system was used to identify the strain as *Sphingomonas parapaucimobilis*, with only 91% identity. Strain identified as *B. zoohelcum* by MALDI-TOF MS was subjected to 16S rRNA gene sequencing analysis to confirm identification by MALDI-TOF MS. Nucleotide alignment using investigated sequences from the GenBank database showed high similarity with *B. zoohelcum* strain. Thus, identification result obtained by MALDI-TOF MS were concordant with those obtained by 16S rRNA gene sequencing analysis.

### phylogenetic analysis

Retrieving the 16S rRNA gene sequences of species showing relatively high similarities to the 16S-D3.13302251, a phylogenetic tree was constructed (Fig. 2). The type strain with the greatest pairwise similarity to strain 16S-D3.13302251 was *B. zoohelcum* ATCC 43767^T^ and *B. zoohelcum* D658^T^, which demonstrated a sequence similarity of 99.71%. BLAST analysis placed the sequence of 16S-D3.13302251 within the group *B. zoohelcum*. In Fig. 2, the strain, 16S-D3.13302251, was clustered with *B. zoohelcum*, and their grouping was supported by bootstrap analysis. Consequently, the phylogenetic tree supported that the strain belonged to species of *B. zoohelcum*.

**FIG 2.**
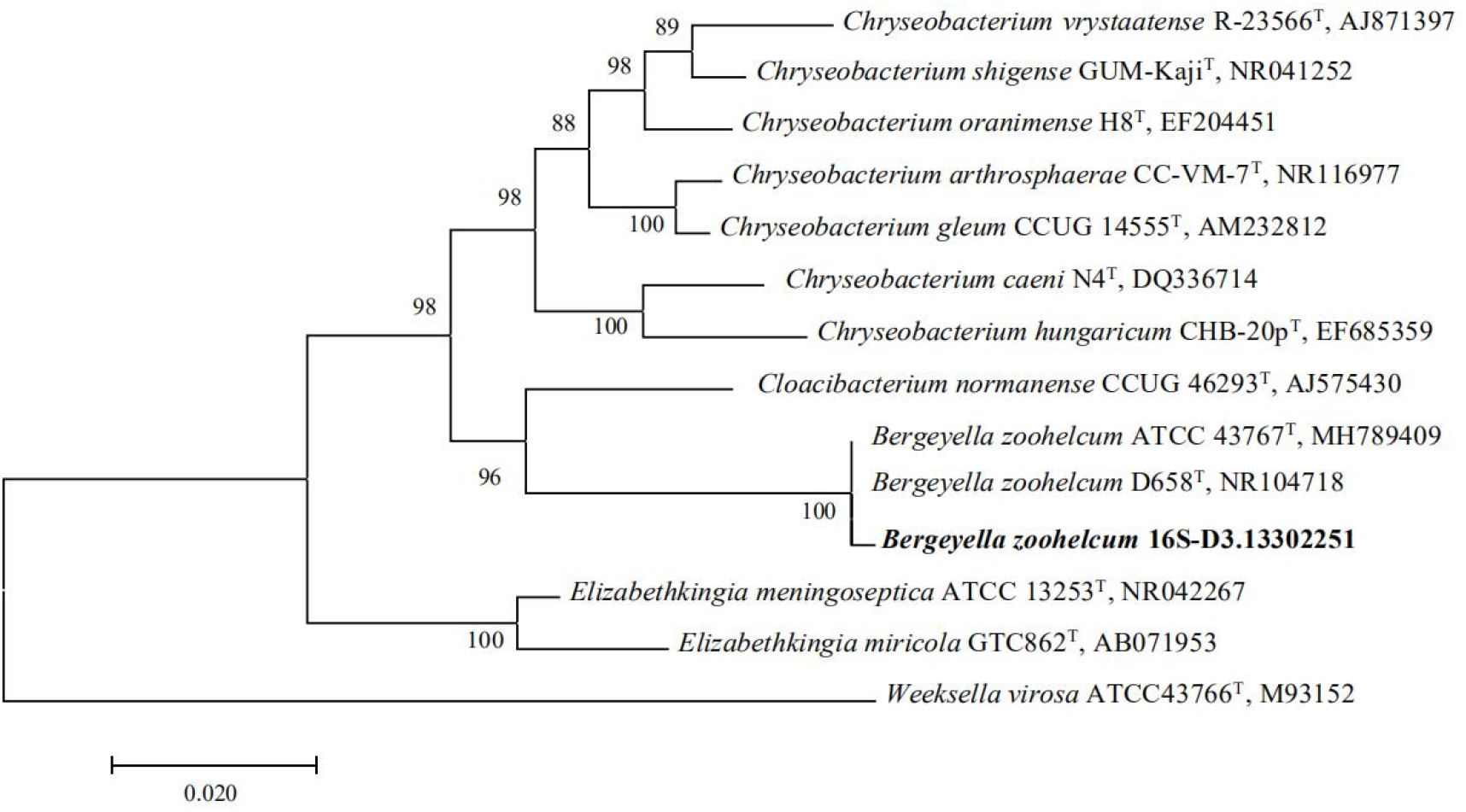
The evolutionary history was inferred using the Neighbor-Joining method. The percentage of replicate trees in which the associated taxa clustered together in the bootstrap test (1000 replicates) are shown next to the branches. The tree is drawn to scale, with branch lengths in the same units as those of the evolutionary distances used to infer the phylogenetic tree. The evolutionary distances were computed using the Maximum Composite Likelihood method and are in the units of the number of base substitutions per site. The analysis involved 14 nucleotide sequences. All positions containing gaps and missing data were eliminated. There were a total of 1271 positions in the final dataset. Evolutionary analyses were conducted in MEGA7.

### Antibiotic susceptibility testing

By now, although there are no accepted CLSI standards for AST or breakpoints for *B. zoohelcum*, the isolate exhibited large inhibition zone (millimeter) for all antimicrobials tested: aztreonam 26 mm, ceftriaxone 30 mm, ciprofloxacin 26 mm, clindamycin 30 mm, erythromycin 25 mm, linezolid 30 mm, teicoplanin 16 mm, trimethoprim-sulfamethoxazole 26 mm, ampicillin 26mm, ampicillin-sulbactam 26mm, vancomycin 17 mm, penicillin 23 mm, cefazolin 27 mm, meropenem26 mm, ertapenem 25 mm, gentamicin 18 mm, levofloxacin 23 mm, cefperazone-sulbactam 28 mm, piperacillin-tazobactam 30 mm, cefazolin 28 mm, cefuroxime 35 mm, which suggested the *B. zoohelcum* isolate was highly susceptible to β-lactams and fluoroquinolones.

## DISCUSSION

*B. zoohelcum* is an uncommon zoonotic pathogen. Historically, this species was referred to as a Centers for Disease Control and Prevention group IIj organism. However, in 1986 Holmes et al. proposed the name *Weeksella zoohelcum* for the group IIj bacteria. *Weeksella virosa* is the only other species in this genus (5). Finally, in 1994 Vandamme et al. proposed a new genus, *Bergeyella*, and renamed *W. zoohelcum B. zoohelcum* based on the genetic differences with *W. virosa* (6). At present, *B. zoohelcum* is the only representative of the genus *Bergeyella*. *B. zoohelcum* is anonfermentative, gram-negative, rod-shaped, non-spore-forming, nonmotile aerobic bacterium. (7)

*B. zoohelcum* has become an increasingly recognized cause of cellulitis, leg abscess, tenosynovitis, septicemia, meningitis and infective endocarditis (8–20), which is closely related to animal bites or contact to dogs, cats or contaminated food (Table 1).

**TABLE 1.**
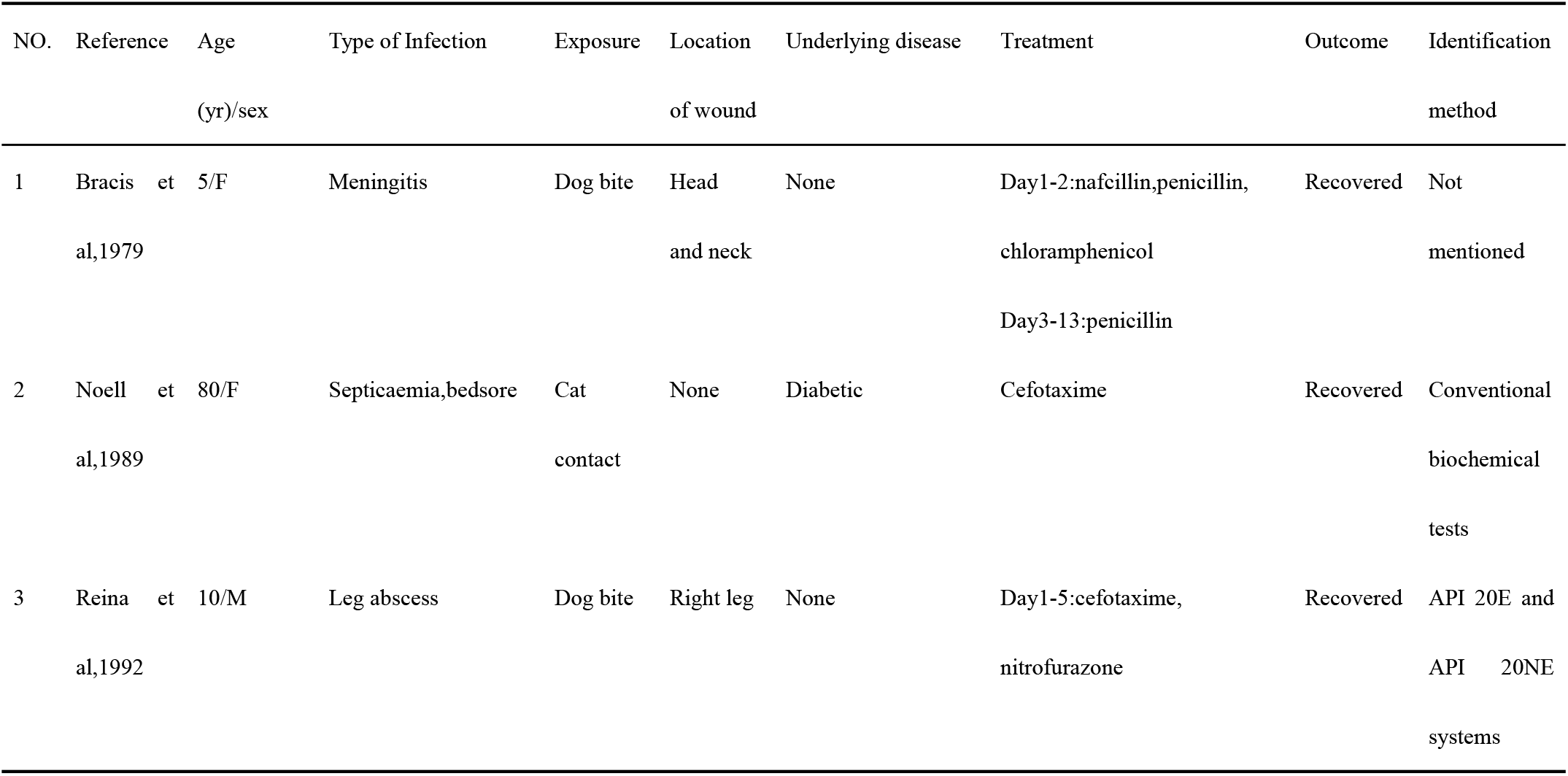
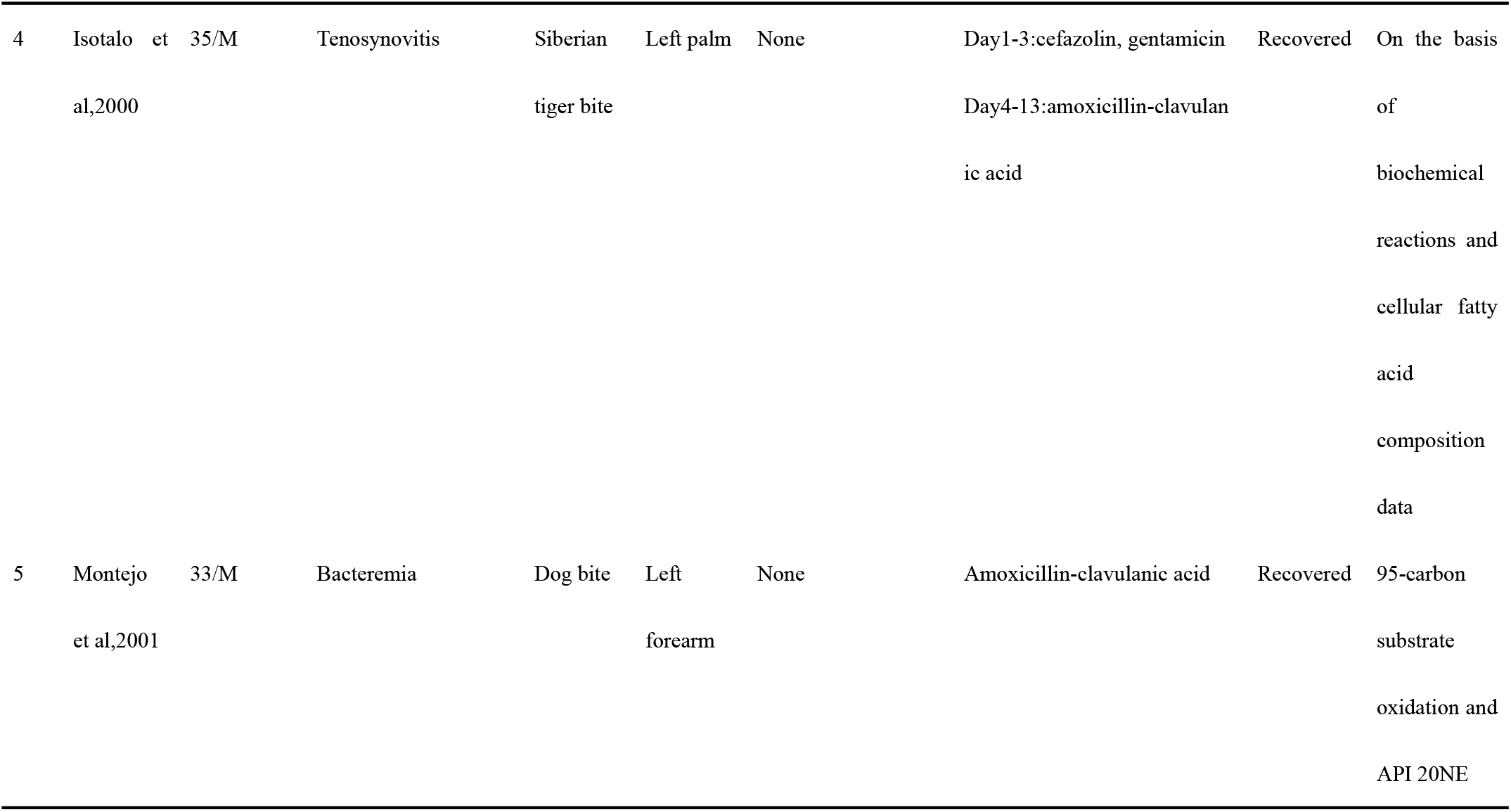
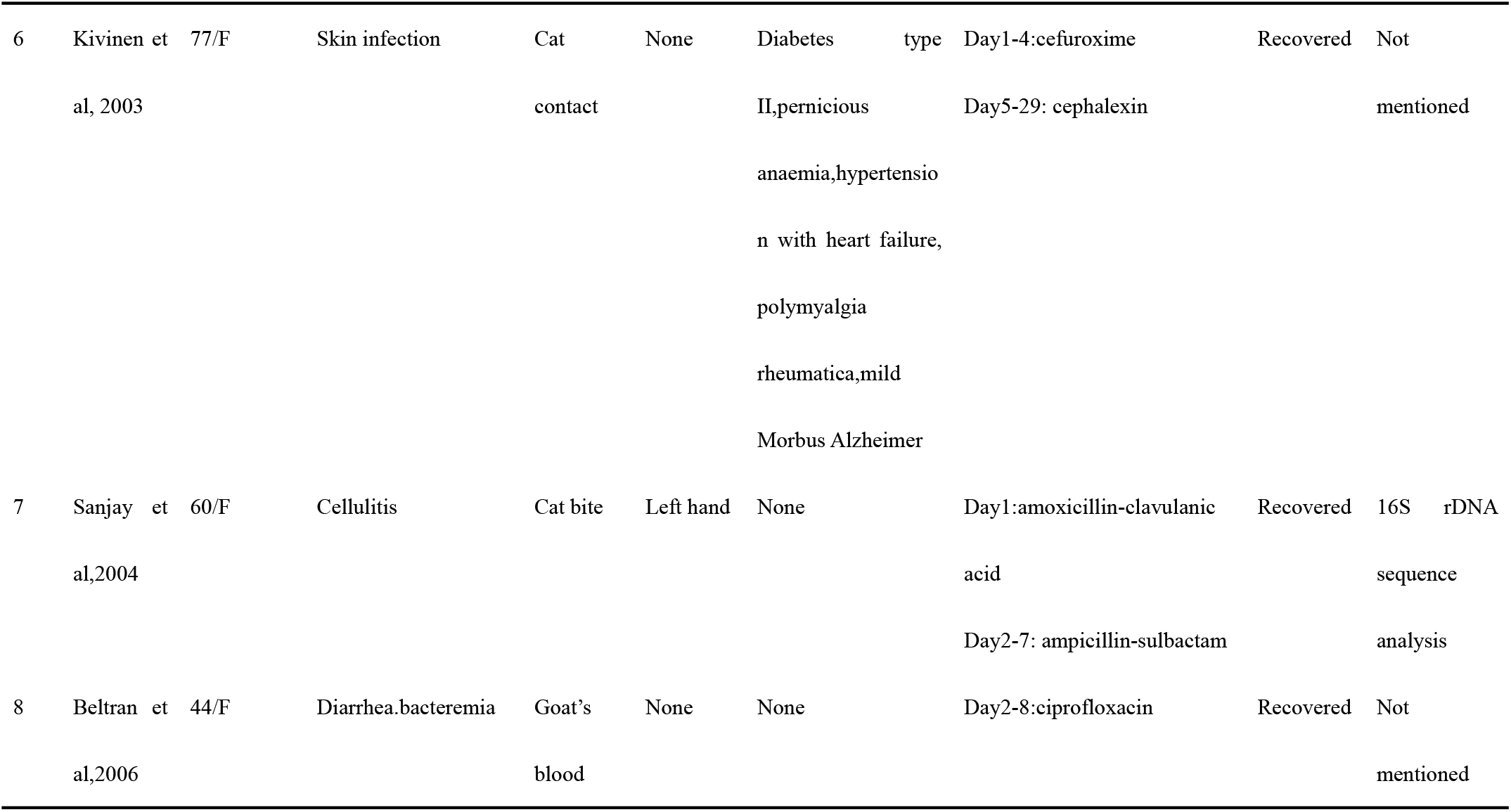
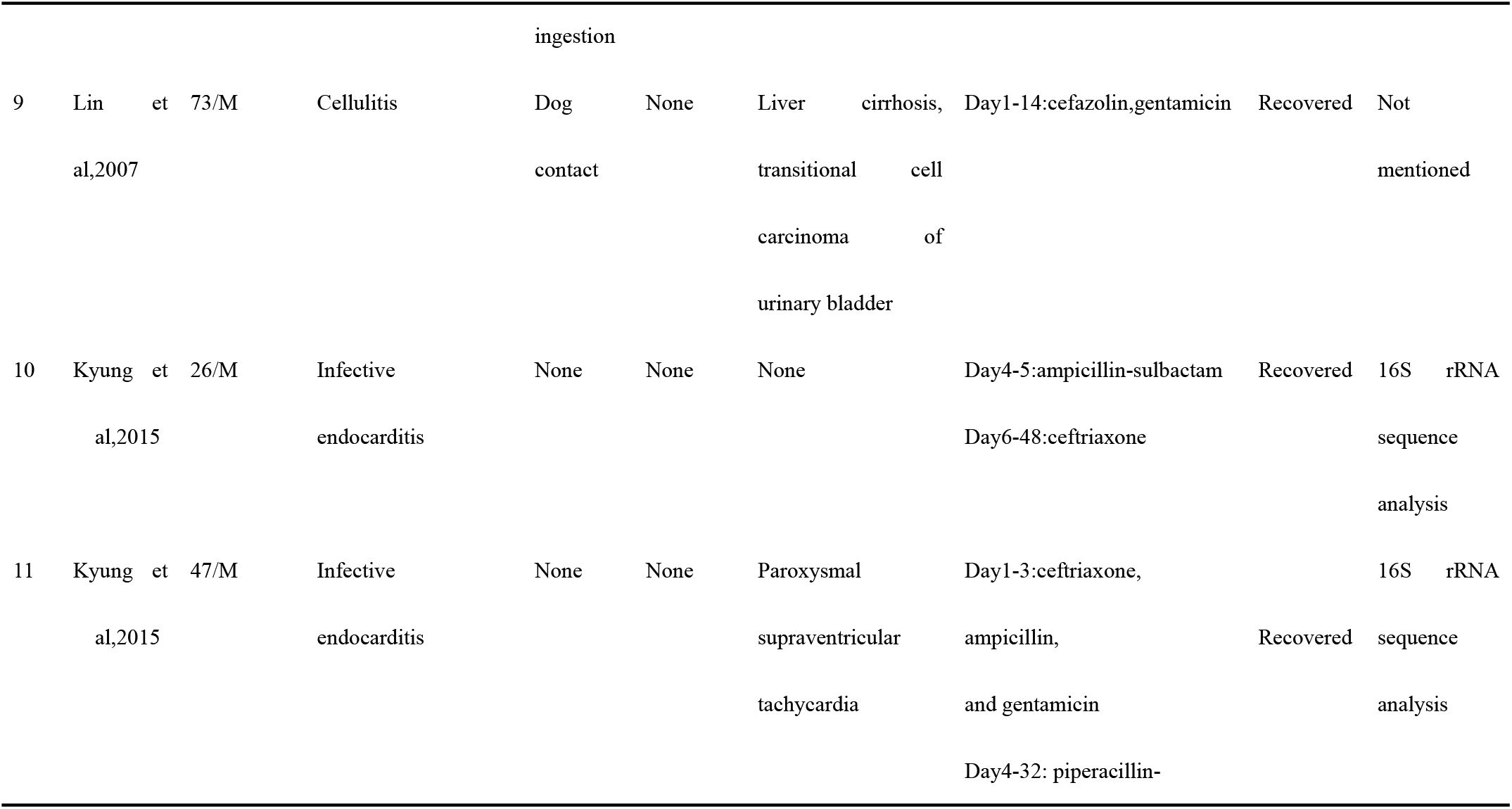
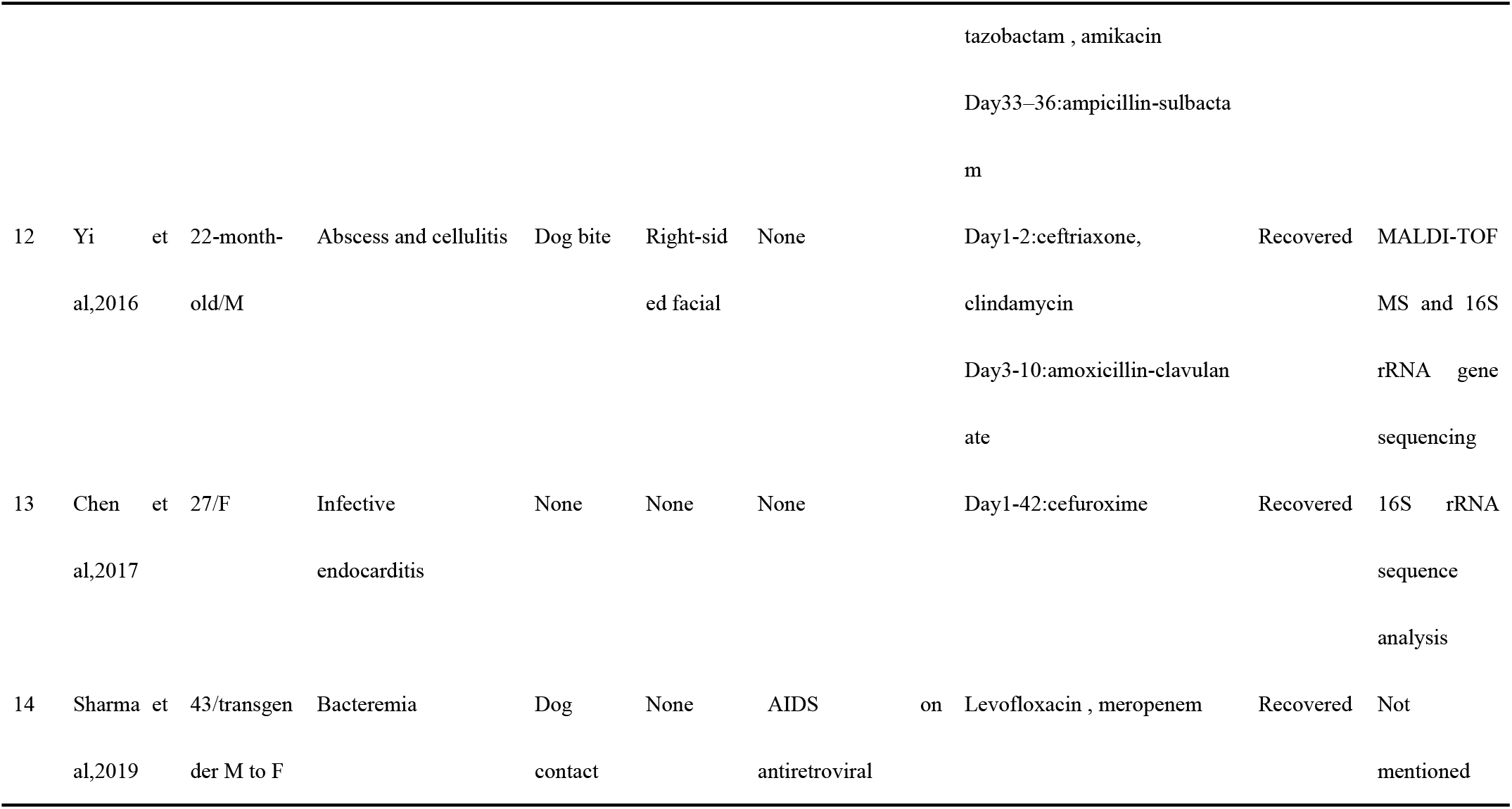
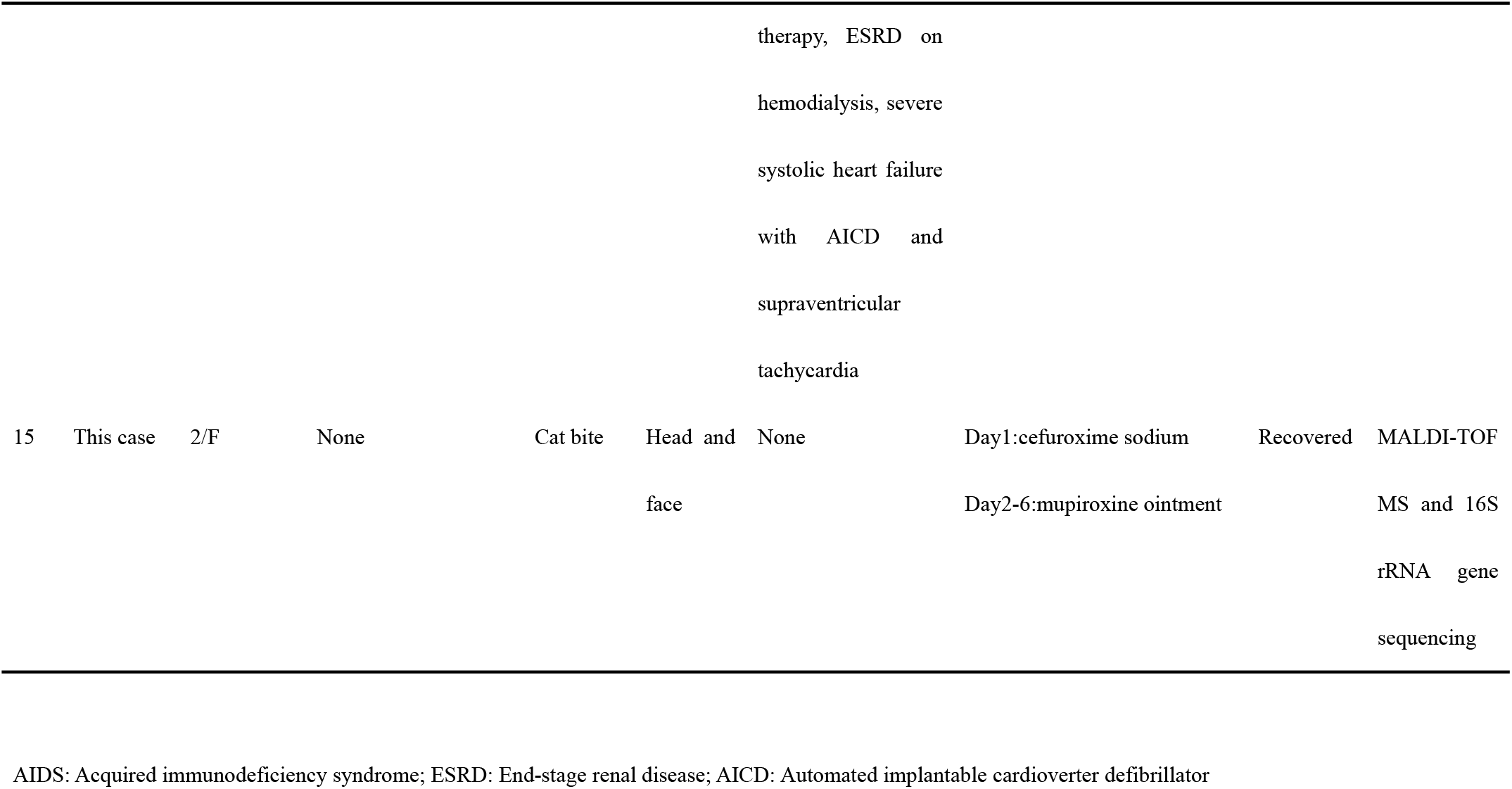
Literature review of *Bergeyella zoohelcum* infections in humans

The reasons for the low detection rate of this strain are as follows: firstly, this strain is a caustic strain and has high requirements for growth environment. Previously reported that light growth of *B. zoohelcum* strain was observed only on the chocolate agar plates after 48 h of incubation (14,18). But it has been reported that after 18 to 24 h incubation the organism grew well on blood and chocolate agar (8,10,17). The strain investigated in our study did not grow on the chocolate plate containing vancomycin. Then, our AST data, indicate that the *B. zoohelcum* strain did not grow around to vancomyci on the Mueller-Hinton agar plates with 5% sheep blood. In addition, the bacteria grew well in the Colombia blood agar plate after cultured for 24 h. Whereas, the strain did not grow on blood plate until 48 hours later. In order to improve the detection rate of bacteria, we recommend using Colombian blood plate and vancomycin-free chocolate plate or delaying the culture time to 48 hours when cultivating bacteria infected by bite infection. Secondly, the *B. zoohelcum* has been misidentified as *Brevundimonas spp*. with the use of an automated identification system (Vitek 2 system) based on conventional phenotypic methodology, or has been considered as a contaminant on culture (8,15,17,19). Initially, our strain was identified by Vitek 2 system as *Sphingomonas parapaucimobilis*.

In the recent years, MALDI-TOF MS is an effective tool for rapid identification of rarely isolated, difficult-to-identify microorganisms, derived from not only human clinical samples but also animal samples. Our strain was identified as *B. zoohelcum* by using MALDI-TOF MS. For a more precise identification, we performed a molecular identification analysis for the stain based on the 16S rRNA gene sequences. Molecular biological characterization has been well-accepted as a powerful tool for identification of bacterial species and strains because of its accuracy. 16S rRNA gene sequencing analysis has shown advantages for identification and differentiation of bacteria from various origins (21). Additionally, the results of phylogenetic analysis also confirmed that the strain belonged to the *B. zoohelcum*. We suggest that when dealing with samples cultured from wound secretions after animal bites, the colony morphology should be carefully observed. If the colony is viscous and difficult to remove from the solid medium, and the oxidase and indole test are positive, we should highly suspect that it is *B. zoohelcum*. It can be seen from table 1 that automated identifification system are not recommended for the identification method of the bacteria. We can identify the organism use the API 20NE systems. It is recommended to use MALDI-TOF MS and sequencing techniques to identify the bacteria.

Whether antibiotics prevent infection after bites remains controversial. Currently, antibiotics are not given routinely, but they are almost always recommended for high-risk wounds, such as deep punctures (particularly if inflicted by cats), those that require surgical repair (22). There is no specific antibiotic treatment recommended for infections caused by *B. zoohelcum*. Nevertheless, it is essential to perform antibiotic susceptibility studies for all clinical isolates until more epidemiological information regarding *B. zoohelcum* is available (10). Optimal selection of empirical antibiotic agents should be based on the most common pathogens. Our patient who was isolated *B. zoolhelcum* from wound, but there was no progressive infection. The reason is attributed to immediate irrigated and debridement of wounds and minimize the risk of the child developing serious or even fatal infections. After being bitten by an animal, adherence to standard principles of wound management provides the best defense against bacterial infections. Copious irrigation at high pressure markedly decreases the concentration of bacteria in contaminated wounds. Debridement of devitalized tissue further decreases the likelihood of infection, with repair only when the possibility of infection has been eliminated (23). Furthermore, we know that local blood supply is excellent in the head, neck, scalp and face, so wounds are at lower risk for infection than other sites (24). Moreover, choose appropriate antibiotics for empirical coverage when indicated for prophylaxis or for treatment of infection. An adequate antibiotic regimen should provide coverage for the potential pathogenic aerobic and anaerobic flora from the mouth of the animal inflicting the bite. However, patients can be treated with agents that have been demonstrated to be effective against strains isolated from animals. Cefotaxime, amoxicillin-clavulanic acid cefuroxime, ampicillin-sulbactam, ciprofloxacin, ceftriaxone, levofloxacin and meropenem have been used to treat patients successfully. Use of amoxicillin-clavulanic acid and ampicillin-sulbactam is appropriate for the treatment of bite-related infection and is a reasonable choice for possible coinfection with other pathogens including *Pasteurella multocida* and anaerobess (Table 2). The little girl was treated with cefuroxime sodium with a good outcome.

**TABLE 2.**
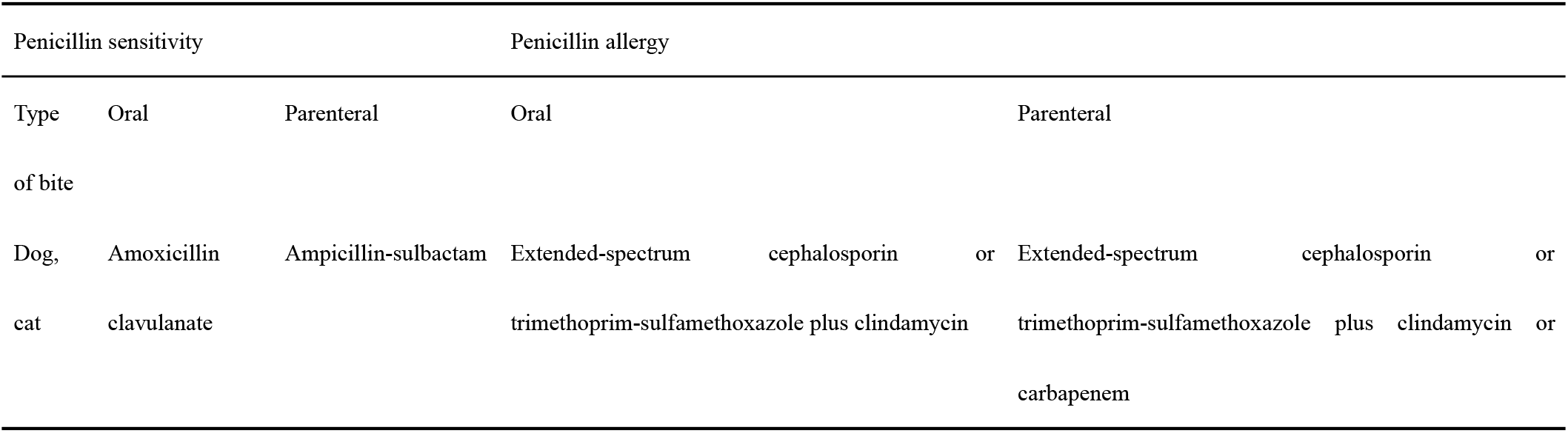
Empirical Antibiotics for Animal Wounds

With our understanding of the fastidous bacteria and the widespread use of MALDI-TOF MS, 16S rRNA gene sequence and next-generation sequences (NGS). We should be able to isolate more *B. zoohelcum* from animal bite wounds and identify the pathogen accurately and quickly.

## ACKNOWLEDGMENTS

We thank Xiao-Qiang Han for assistance in organism identification of 16S rRNA sequence analysis..

We declare no conflict of interests.

## REFERENCES

1. Griego RD, Rosen T, Orengo IF, Wolf JE. Dog, cat, and human bites: a review. J Am Acad Dermatol. 1995 Dec;33(6):1019–1029.

2. Oehler RL, Velez AP, Mizrachi M, Lamarche J, Gompf S. Bite-related and septic syndromes caused by cats and dogs. Lancet Infect Dis. 2009 Jul;9(7):439–447.

3. Chun YT, Berkelhamer JE, Herold TE. Dog bites in children less than 4 years old. Pediatrics. 1982 Jan;69(1):119–120

4. Talan DA, Citron DM, Abrahamian FM, Moran GJ, Goldstein EJ. Bacteriologic analysis of infected dog and cat bites. Emergency Medicine Animal Bite Infection Study Group. N Engl J Med. 1999 Jan 14;340(2):85–92.

5. Holmes B, Steigerwalt AG, Weaver RE, Brenner DJ. 1986. *Weeksella zoohelcum* sp. nov. (formerly group IIj) from human clinical specimens. Syst. Appl. Microbiol. 8:191–196.

6. Vandamme P, Bernardet JF, Segers P, Kersters K, Holmes B. 1994.New perspectives in the classification of the flavobacteria: description of Chryseobacterium gen. nov., Bergeyella gen. nov., and Empedobacter nom. rev.Int. J. Syst. Bacteriol. 44:827–831.

7. Steinberg JP, Del Rio C. Other Gram-negative and Gram variable bacilli-*Weeksella* and *Bergeyella*. In: Mandell GL, Bennett JE, Dolin R (eds). Mandell, Douglas, and Bennett’s Principles and Practice of Infectious Diseases, 6th edition. Philadelphia: Elsevier, 2005:2762.

8. Bracis R, Seibers K, Julien RM. Meningitis caused by group II J following a dog bite. West J Med. 1979 Nov;131(5):438–440.

9. Noell F, Gorce MF, Garde C, Bizet C. Isolation of *Weeksella zoohelcum* in septicaemia. Lancet. 1989 Aug 5;2(8658):332.

10. Reina J, Borrell N. Leg abscess caused by *Weeksella zoohelcum* following a dog bite. Clin Infect Dis. 1992 May;14(5):1162–1163.

11. Isotalo PA, Edgar D, Toye B. Polymicrobial tenosynovitis with *Pasteurella multocida* and other gram negative bacilli after a Siberian tiger bite. J Clin Pathol. 2000 Nov;53(11):871–872.

12. Montejo M, Aguirrebengoa K, Ugalde J, Lopez L, Saez Nieto JA, Hernández JL. *Bergeyella zoohelcum* bacteremia after a dog bite. Clin Infect Dis. 2001 Nov 1;33(9):1608–1609.

13. Kivinen PK, Lahtinen MR, Ruotsalainen E, Harvima IT, Katila ML. *Bergeyella zoohelcum* septicaemia of a patient suffering from severe skin infection. Acta Derm Venereol. 2003;83(1):74–75.

14. Shukla SK, Paustian DL, Stockwell PJ, Morey RE, Jordan JG, Levett PN, Frank DN, Reed KD. Isolation of a fastidious *Bergeyella* species associated with cellulitis after a cat bite and a phylogenetic comparison with *Bergeyella zoohelcum* strains. J Clin Microbiol. 2004 Jan;42(1):290–293.

15. Beltran A, Bdiiwi S, Jani J, Recco RA, Go EE, Zaman MM. A case of *Bergeyella zoohelcum* bacteremia after ingestion of a dish prepared with goat blood. Clin Infect Dis. 2006 Mar 15;42(6):891–892.

16. Lin WR, Chen YS, Liu YC. Cellulitis and bacteremia caused by *Bergeyella zoohelcum*. J Formos Med Assoc. 2007 Jul;106(7):573–576.

17. Sohn KM, Huh K, Baek JY, Kim YS, Kang CI, Peck KR, Lee NY, Song JH, Ko KS, Chung DR. A new causative bacteria of infective endocarditis, *Bergeyella* cardium sp. nov. Diagn Microbiol Infect Dis. 2015 Mar;81(3):213–216.

18. Yi J, Humphries R, Doerr L, Jerris RC, Westblade LF. *Bergeyella zoohelcum* Associated with Abscess and Cellulitis After a Dog Bite. Pediatr Infect Dis J. 2016 Feb;35(2):214–216.

19. Chen Y, Liao K, Ai L, Guo P, Huang H, Wu Z, Liu M. Bacteremia caused by *Bergeyella zoohelcum* in an infective endocarditis patient: case report and review of literature. BMC Infect Dis. 2017 Apr 12;17(1):271.

20. Sharma S, Salazar H, Sharma S, Nasser MF, Dahdouh M. *Bergeyella zoohelcum* Bacteremia from Therapy Dog Kisses. Cureus. 2019 Apr 18;11(4):e4494.

21. Muramatsu Y, Haraya N, Horie K, Uchida L, Kooriyama T, Suzuki A, Horiuchi M. *Bergeyella zoohelcum* isolated from oral cavities of therapy dogs. Zoonoses Public Health. 2019 Dec;66(8):936–942.

22. Fleisher GR. The management of bite wounds. N Engl J Med. 1999 Jan 14;340(2):138–140.

23. Javaid M, Feldberg L, Gipson M. Primary repair of dog bites to the face: 40 cases. J R Soc Med. 1998 Aug;91(8):414–416.

24. Guy RJ, Zook EG. Successful treatment of acute head and neck dog bite wounds without antibiotics. Ann Plast Surg. 1986 Jul;17(1):45–48.

